# X-ray imaging of 30 year old wine grape wood reveals cumulative impacts of rootstocks on scion secondary growth and harvest index

**DOI:** 10.1101/2022.05.17.492371

**Authors:** Zoë Migicovsky, Michelle Y. Quigley, Joey Mullins, Tahira Ali, Joel F. Swift, Anita Rose Agasaveeran, Joseph D. Dougherty, Brendan Michael Grant, Ilayda Korkmaz, Maneesh Reddy Malpeddi, Emily L. McNichol, Andrew W. Sharp, Jackie L. Harris, Danielle R. Hopkins, Lindsay M. Jordan, Misha T. Kwasniewski, R. Keith Striegler, Asia L. Dowtin, Stephanie Stotts, Peter Cousins, Daniel H. Chitwood

## Abstract

- Annual rings from 30 year old vines in a California rootstock trial were measured to determine the effects of 15 different rootstocks on Chardonnay and Cabernet Sauvignon scions. Viticultural traits measuring vegetative growth, yield, berry quality, and nutrient uptake were collected at the beginning and end of the lifetime of the vineyard.
- X-ray Computed Tomography (CT) was used to measure ring widths in 103 vines. Ring width was modeled as a function of ring number using a negative exponential model. Early and late wood ring widths, cambium width, and scion trunk radius were correlated with 27 traits.
- Modeling of annual ring width shows that scions alter the width of the first rings but that rootstocks alter the decay thereafter, consistently shortening ring width throughout the lifetime of the vine. The ratio of yield to vegetative growth, juice pH, photosynthetic assimilation and transpiration rates, and stomatal conductance are correlated with scion trunk radius.
- Rootstocks modulate secondary growth over years, altering hydraulic conductance, physiology, and agronomic traits. Rootstocks act in similar but distinct ways from climate to modulate ring width, which borrowing techniques from dendrochronology, can be used to monitor both genetic and environmental effects in woody perennial crop species.

## Introduction

Grafting is the joining of plant tissues together, through either natural or artificial means (Gaut *et al.*, 2019). When a root system (the rootstock) is grafted to a shoot system (the scion, pronounced *sai* • uhn), a graft junction is formed, with vascular connections connecting the two systems into a single organism (Thomas & Frank, 2019). Hormones, tissue regeneration, and molecular pathways regulating vascular development are important to this process (Melnyk, 2017; Nanda & Melnyk, 2018). Often, the rootstock and scion are genetically distinct, combining two genotypes into a single chimera if no genetic incompatibility arises (Thomas *et al.*, 2022). When genetically distinct root and shoot systems are grafted together (a heterograft), they can be compared to different grafted rootstock and scion genotypes, the same genotype grafted to itself (a homograft), or an ungrafted plant (own-rooted) (Frank & Chitwood, 2016). Any difference in scion or rootstock traits relative to a different combination of grafted genotypes can be used to infer the genetic basis of reciprocal communication between root and shoot systems. Graft-induced signaling between genotypes has been used extensively during domestication, breeding, and agricultural improvement of crops (Williams *et al.*, 2021).

Grafting can be performed across the flowering plants (Reeves *et al.*, 2022). Especially in long-lived, woody perennials, grafting can be necessary for cultivation and even domestication itself (Warschefsky *et al.*, 2016). In perennial species, grafting has been used to modulate scion architecture through dwarfing, to alter fruit bearing through precocity and productivity, to change fruit quality, and to confer pathogen resistance and abiotic stress tolerance (Warschefsky *et al.*, 2016). Over 80% of vineyards grow grafted grapes, a process that became widespread in response to the phylloxera *(Daktulosphaira vitifoliae* Fitch) crisis, in which grafting to tolerant and/or resistant rootstocks derived from North American *Vitis* species allowed wine and table grape scions to grow in infested soils (Ollat *et al.*, 2016). Subsequently, grapevine rootstocks were recognized as not only conferring resistance to pathogens besides phylloxera (Cousins & Walker, 2002; Hwang *et al.*, 2010; Ferris *et al.*, 2012), but also salinity and drought tolerance (Zhang *et al.*, 2002; Serra *et al.*, 2014), as well as altering scion mineral composition (Walker *et al.*, 2004; Gautier *et al.*, 2018, 2020; Migicovsky *et al.*, 2019; Harris *et al.*, 2022) and the chemistry and maturation of berries (Ruhl, 1989; Walker *et al.*, 2000; Kodur, 2011; Cheng *et al.*, 2017). One of the most desirable properties that grapevine rootstocks can affect is Ravaz (or harvest) index: the ratio of yield to pruning weight (or, 1-year-old cuttings weighed the following dormant season at pruning). A high Ravaz index is desirable to maximize yield and minimize vine management. However, in some cases, a Ravaz index may be too high, indicating that the vine is overcropped and often resulting in reduced fruit quality and reduced vine size (Bravdo *et al.*, 1984). Grapevine rootstocks modulate both components of Ravaz index, yield and vigor (McCarthy & Cirami, 1990; Ezzahouani & Williams, 1995; Jones *et al.*, 2009; Migicovsky *et al.*, 2021). Water and nutrient uptake are two of the principal means by which rootstocks affect reproductive and vegetative growth, consequently also influencing berry composition (Keller, 2020).

Although rootstocks can influence grapevine growth, yield, and berry qualities, the primary source of variation in these traits is the environment (Kidman *et al.*, 2014; Keller, 2020). In non-grafted plants, the root and shoot systems are in constant communication with each other. In particular, the root and shoot systems must communicate to enable hydraulic conductivity, the movement of water from the roots through the vasculature of plants to the leaves and other aerial parts where it is transpired (evaporated) or guttated (exuded). Through water uptake and vasculature, both environment and rootstocks converge on an often overlooked component of plant growth in perennial crops: secondary growth. The effects of the environment on secondary growth in woody perennials can be so strong that, using methods from dendrochronology, the widths of tree rings can be used to infer water availability and the length of the growing season. In angiosperms, the vascular cambium divides during the growing season to create xylem vessels and fibers (Rathgeber *et al.*, 2016). The daughter cells enlarge, thicken secondary walls, and eventually undergo cell death to form the empty lumen that transports water from the root to shoot systems. Grapevines and other vines and lianas have ring porous wood and produce wider vessels early in the season and narrow vessels later (Pratt, 1974; Ewers *et al.*, 1990; Wheeler & LaPasha, 1994). Grapevine annual ring width is environmentally responsive. In 14 year old *Vitis vinifera* L. cv. Merlot grafted to 140 Ruggeri (Munitz *et al.*, 2018) and 12 year old *V. vinifera* L. cv. ‘Cabernet sauvignon’ grafted to 140 Ruggeri (Netzer *et al.*, 2019), annual ring width, vessel diameter, and hydraulic conductivity all increase with applied water, most strongly at the beginning of the growing season when cambial activity is strongest (Berstein & Fahn, 1960).

Grapevine wood anatomy is also affected by grafting. Five year old *V. vinifera* L. cv. Piedirosso vines grafted to 420A have more numerous, narrower vessels in late wood compared to ungrafted counterparts, conferring safer water transport under drought conditions (De Micco *et al.*, 2018). A comparison of wood anatomy between seven year old *V. vinifera* L. cv. Cabernet Sauvignon vines grafted to Riparia Gloire, 420A, and 1103 Paulsen found that a narrow rootstock stem size restricted hydraulic conductivity and affected physiological performance compared to a smoother graft junction with less discrepancy in rootstock and scion stem diameters (Shtein *et al.*, 2017). The extent and success of grafting may also play a role in the effect of rootstock on scion. For example, in a recent study of the effects of alignment of scion and rootstock, vine growth was significantly impacted in the nursery and the first year of establishment when there was partial alignment. However, in years 2 and 3, these differences disappeared (Marín *et al.*, 2022).

Here, using X-ray Computed Tomography (CT) to examine ring widths in 30 year old Chardonnay and Cabernet Sauvignon scion wood grafted to 15 different rootstocks from a California vineyard, we measured the effects of rootstocks on secondary growth. Within the size ranges of each scion, rootstock additively modulates scion trunk radius across a continuous range differing by up to 143% in width. Modeling ring width as a function of ring number, we find that scion modulates the widths of the first rings, whereas rootstock modulates decay, cumulatively affecting secondary growth throughout the lifetime of the vine. Traits collected early and late during the lifetime of the vineyard show correlations with scion trunk radius, including juice pH and Ravaz index, reflecting effects on vegetative growth. Consistent with previous work, scion trunk radius is also correlated with physiological performance, affecting photosynthetic assimilation, transpiration rate, and stomatal conductance. Our results show that the cumulative effects of grapevine rootstocks on viticultural traits act consistently over decades by altering physiological performance through scion wood anatomy.

## Materials and Methods

### Vineyard history, management, and design

A complete history of the vineyard sampled in this study is available in Migicovsky *et al.* (2021), reprinted here for convenience. In 1991 rootstocks were planted near Lodi in San Joaquin County, California, before scionwood was whip-grafted to the planted rootstock in 1992. The scion cutting was a cane (1 year old wood) when it was grafted. The soil type was a Tokay fine sandy loam soil. Vines were grafted to the following rootstocks: Freedom, Ramsey, 1103 Paulsen, 775 Paulsen, 110 Richter, 3309 Couderc, Kober 5BB, SO4, Teleki 5C, 101-14 Mgt, 039-16, 140 Ruggeri, Schwarzman, 420 A, and K51-32. The parental *Vitis* species for each rootstock are presented in **Table 1** (Hardie & Cirami, 1988; Riaz *et al.*, 2019). The two scion varieties were Chardonnay (selection FPS 04) and Cabernet Sauvignon (selection FPS 07). Rows were oriented east-west with vine spacing of 2.13 m by 3.05 m. The trellis system was a bilateral cordon with fixed foliage wires and the vines were cordon trained and spur pruned. The experimental design was a randomized complete block design, split between Chardonnay and Cabernet Sauvignon. There were four replications per treatment (rootstock). There were eight or nine vines per plot, except for Kober 5BB and SO4, which had four or five vines each, to fit all treatments in the block.

**Table 1:**
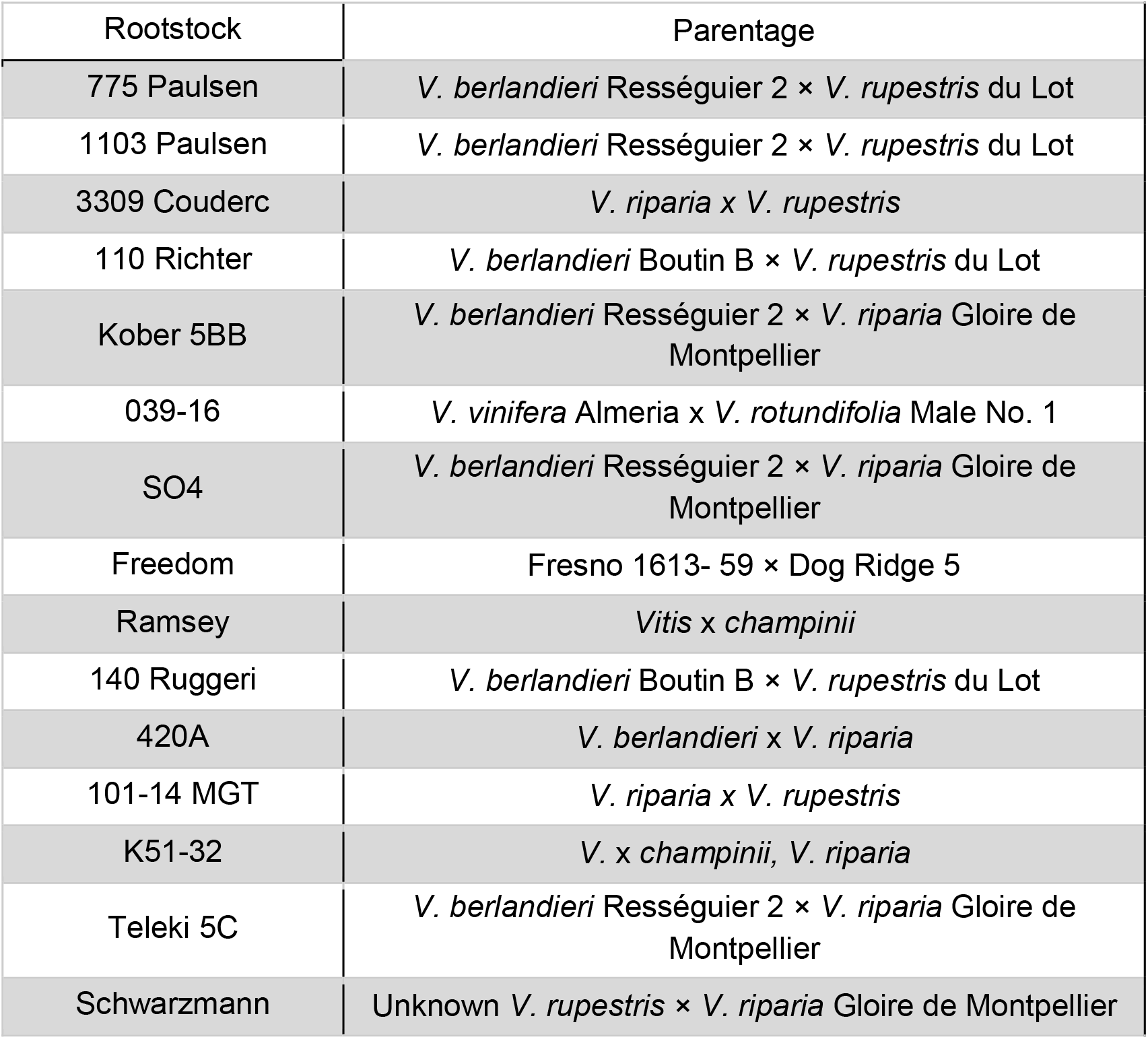
Rootstock parentage.

### Trait collection

Historical data across 8 traits and all 15 rootstocks were collected for 5 years, from 1995 to 1999, and described in Migicovsky *et al.* (2021). These traits were: soluble solids content, titratable acidity, pH, berry weight, cluster number, yield, pruning weight, and Razav index (a measurement of crop load calculated by dividing yield by pruning weight from the following dormant season). Contemporary data for all 15 rootstocks were also collected for 4 years, from 2017 to 2020, for the 8 traits included in the historical data set. Berry measurements were not calculated for 5BB Kober and SO4 in 2017 only, due to sample mislabelling at the time. In addition to the 8 traits from the historical data, numerous additional traits were collected for one or more years in the contemporary data set, including petiole nutrition measurements, cordon length, and yeast assimilable nitrogen (YAN). For a full summary of the number of measured samples for each trait included in analysis, for each scion, for each year, see **Supporting Information Table S1**.

In addition to these datasets, sampling took place at the vineyard for physiological measurements in 2018 and 2019. This sampling was performed using only 3 of the rootstocks, for each scion: Teleki 5C, Freedom, and 1103 Paulsen. In 2018, sampling was performed once a week for 8 weeks from June 19th, 2018 to August 6th, 2018. In 2019, sampling was performed once a week for 7 weeks from June 17th, 2019 to July 29th, 2019. For each rootstock x scion combination, 3 vines were sampled, and for each vine, 2 leaves were measured. Measurements were subsequently averaged across leaves from a particular vine. The same 2 leaves were measured for both physiological traits and leaf temperature.

Physiological traits were measured using a portable photosynthesis system (LI-6800, LICOR Biosciences, Lincoln, NE, USA) on clear sky days. Flow (600 μmol s^−1^), H_2_O (RH_air 50%), C_2_O (CO2_r 400 μmol mol^−1^), temperature (Tleaf 33 °C), and light (1800 μmol m□^−2^s^−1^) were kept constant throughout sampling periods. Three measurements were taken: stomatal conductance (GSW, mol m□^2^s□^1^), photosynthetic CO2 rate (A, μmol m□^2^□s□^1^), and transpiration rate (E, mol m ^2^ s□^1^). Fully expanded, sunlight leaves were measured, and each leaf was from a different shoot on the vine and selected to represent the canopy as a whole. Measurements were taken midday (approximately 10:30 am – 2:30 pm). Leaf temperature measurements were taken either immediately before or after the physiological measurements using an infrared thermometer (Extech 42515), scanning the same leaves measured for physiological traits.

### X-ray Computed Tomography and ring measurement

All trunk samples were collected on March 2, 2021 using a chainsaw. Trunk sections were roughly 20 cm long and taken 15-20 cm below the cordon wire (101.6 cm height) at the head of the vine, normally just below cordon split. Given that 1 year old scionwood was grafted in 1992 and cuttings of it were taken in 2021, the physiological age of the trunk samples was 30 years old, although the vineyard itself was only 29 years old. Samples were placed in polyethylene bags, sealed, and shipped to Michigan State University. CT images were taken from the middle of the vine samples. The total image height was approximately 95 cm. Images were scanned at 75 kV and 450 μamps in continuous mode with 2880 projections and 2 frames averaged on a North Star Imaging X3000. The focal spot size was 33.75 microns and the detector was set to 12.5 frames per second. Scan time was 8 minutes. Scans were reconstructed using the efX-CT software from North Star Imaging (Rogers, MN). Final voxel resolution was 63.9 μm and pixel resolution was 63.9 μm by 63.9 μm (0.00639 cm by 0.00639 cm). From each 3D CT image, three individual slices were taken for analysis. One slice was from the top of the image, one was from the middle of the image, and the last was from the bottom of the image. All slices were aligned to be taken perpendicular to the vine samples. In ImageJ (Abràmoff *et al.*, 2004). A thin line was drawn from the center of the pith to the bark avoiding eccentric growth and crossing rings as perpendicular as possible. Landmarks were placed according to **Figure 1A**. Euclidean distance converting from pixels to centimeters was used to calculate ring widths. Models of ring width as a function of ring number were fitted with the three slices per each vine. Ring width measurements across the 3 slices for each vine were averaged for trait correlation analysis.

**Figure 1:**
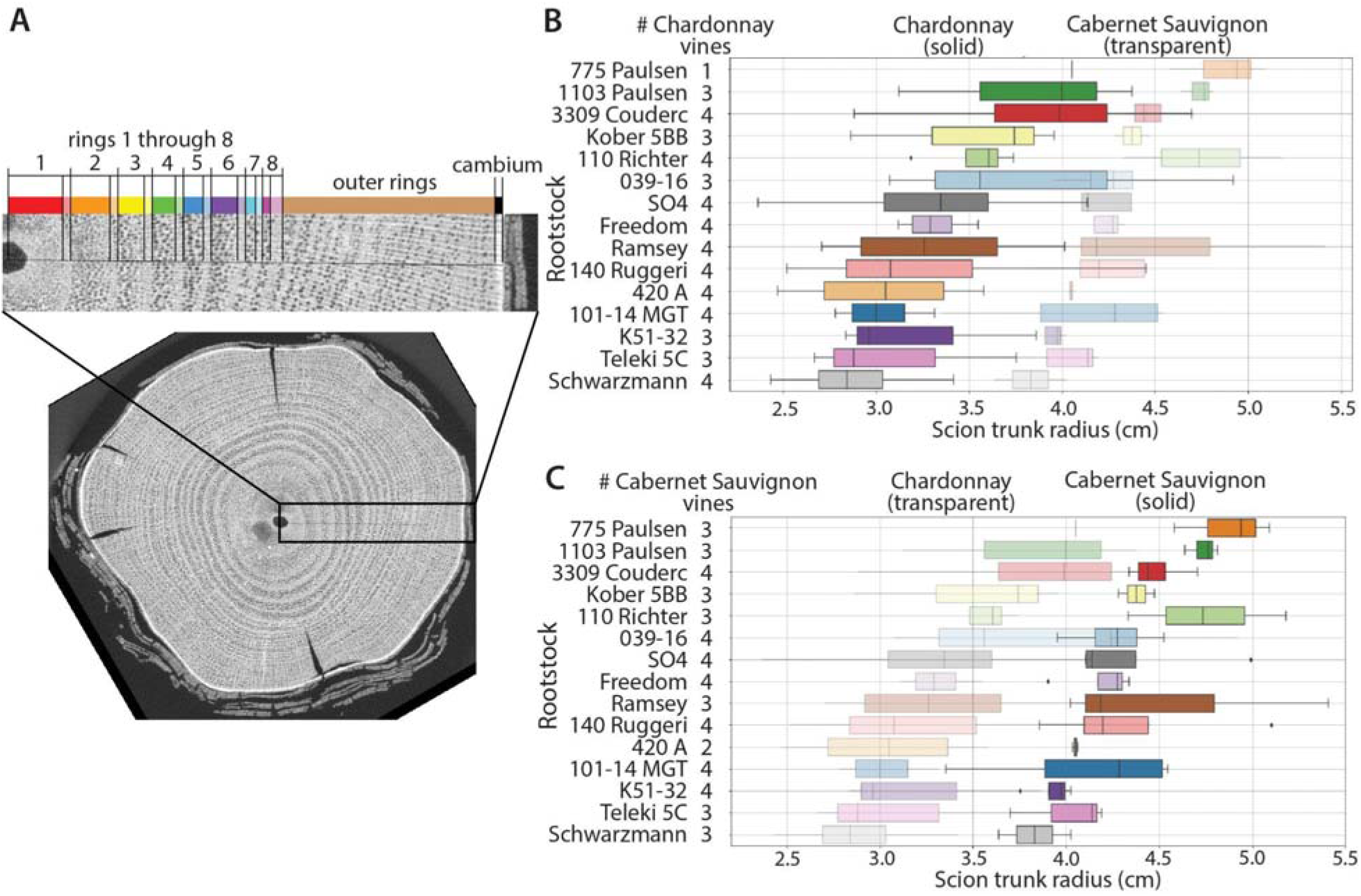
Grapevine rings and scion trunk radii. **A)** X-ray CT cross-section through a grapevine trunk. Along a line from the center of the pith to bark, landmarks are placed indicating early (darker shade color) and late (lighter shade color) wood, the remaining outer rings, and cambium. **B-C)** Boxplots showing distributions of scion trunk radii (cm) in **B)** Chardonnay (solid) relative to Cabernet Sauvignon (transparent) and number of Chardonnay vines measured and **C)** Cabernet Sauvignon (solid) relative to Chardonnay (transparent) and number of Cabernet Sauvignon vines measured.

### Data analysis

Data analysis was performed in Python (v. 3.7.3) using Jupyter notebooks (v. 1.0.0) (Kluyver *et al.*, 2016). Numpy (v. 1.19.4) (Harris *et al.*, 2020) and Pandas (v. 1.3.5) (McKinney, 2011) were used to work with arrays and dataframes and Matplotlib (v. 3.1.0) (Hunter, 2007) and Seaborn (v. 0.11.2) (Waskom, 2021) were used for visualization. The negative exponential model *ring width*. = *A* + *B* * e-^*k*\**ringnumber*^ was fit using the curve_fit() function from Scipy (v. 1.7.3) (Virtanen *et al.*, 2020). The Statsmodels module (v. 0.13.2) (Seabold & Perktold, 2010) was used to perform ordinary least squares (OLS) and analysis of variance (ANOVA) modeling as well as for using the Benjamini-Hochberg (BH) procedure for multiple-test correction. The .corr() function in Pandas was used to calculate Spearman’s rank correlation coefficient. For correlations between scion trunk radius and physiological traits, the repeated measures correlation coefficient (r_rm_) was calculated using the Pingouin module (v. 0.5.1) (Vallat, 2018). The repeated measures correlation coefficient (Bakdash & Marusich, 2017) fits an overall correlation coefficient on a population of repeated measures. In our case the repeated measures are physiological measurements on scions with three different rootstocks (Teleki 5C, Freedom, and 1103 Paulsen) on 15 different dates across 2018 and 2019. All code and data to reproduce the results and visualizations in this manuscript, with comments and narrative, can be found in a Jupyter notebook https://github.com/DanChitwood/grapevine_rings/blob/main/grapevine_ring_analysis_v2.ipynb.

## Results

### Variation in scion trunk radius arises from cumulative effects of rootstocks on ring width

Scion trunk radius varies most strongly by scion, but within scion, rootstock exerts a strong additive effect (**Figure 1B-C**). For the model *scion trunk radius* ~ *scion* * *rootstock*, scion explains 46.53% of variation (p = 3.50×10^−15^) in scion trunk radius and rootstock 16.57% (p = 0.0057), while the interaction effect only 2.42% (p = 0.980) (**Table 2**). Scion shifts the range of scion trunk radii conferred by rootstocks in an additive fashion. Within Chardonnay, the median scion trunk radius arising from the rootstock conferring the widest radius (775 Paulsen, 4.05 cm) is 143% wider than the rootstock conferring the narrowest radius (Schwarzmann, 2.84 cm) (**Figure 1B**). The same rootstocks define the range of scion trunk radii within Cabernet Sauvignon and closely follow the order of that in Chardonnay, with the rootstock conferring the widest radius (775 Paulsen, 4.94 cm) 129% wider than that conferring the narrowest (Schwarzmann, 3.83 cm) (**Figure 1C**).

**Table 2:** **For traits scion trunk radius and** *A, B*, **and** *k* **values from the model** *ring width* = *A* + *B* * *e*-^*k*\**ringnumber*^, **the percent variation explained and p values for each factor in the model *trait* ~ *scion* * *rootstock***

|  | Scion |  | Rootstock |  | Scion x Rootstock |  |
| --- | --- | --- | --- | --- | --- | --- |
| Trait | Percent variation | p value | Percent variation | p value | Percent variation | p value |
| Scion trunk radius | 46.53% | $3.50 \times 10^{-15}$ | 16.57% | 0.0057 | 2.42% | 0.980 |
| $A$ | 1.78% | 0.160 | 22.21% | 0.0565 | 10.35% | 0.632 |
| $B$ | 13.67% | 0.000068 | 19.01% | 0.0593 | 10.64% | 0.470 |
| $k$ | 0.33% | 0.527 | 22.68% | 0.0298 | 16.86% | 0.139 |

Tree ring widths can be modeled as a function of ring number using a negative exponential model (Fritts *et al.*, 1969). We applied the model *ring width* = *A* + *B* * *e*-^*k*\**ring number*^ to measure how scion and rootstock modulate ring width across the scion trunk. Comparing the overall model to models fitted with +/− 1.5 standard deviations from the mean values for *A, B*, and *k* shows how each of these variables acts (**Figure 2A**). *A* transposes ring width, including the asymptote, up and down across all rings, *B* affects the widths of the first rings but not the asymptote of the later rings, and *k* affects the decay, how rapidly the ring width approaches the asymptote. For all rings, modeled width was higher in Cabernet Sauvignon than Chardonnay (**Figure 2B-C**), consistent with scion trunk radii (**Figure 1**). If models of ring width are calculated for each rootstock for each scion, differences in decay are observed by rootstock (**Figure 2D-E**). For example, one of the rootstocks conferring a wider scion trunk radius (1103 Paulsen, dark green in **Figure 2D-E**) maintains higher ring widths, whereas one of the rootstocks conferring a narrower scion trunk radius (Teleki 5C, dark pink in **Figure 2D-E**) rapidly drops in ring width value after the first couple rings. Additionally Cabernet Sauvignon has higher ring widths in the first couple rings compared to Chardonnay, but both scions fall to a similar asymptote around 0.2 cm in later rings. To quantify these trends, we calculated modeled *A, B*, and *k* values for each vine (**Figure 2F-H**) and analyzed how they vary across scion and rootstock variables using the model ***trait*** ~ ***scion*** * ***rootstock*** (**Table 2**). No terms were significant for *A*, which is the translation of the curve along the *y* axis, but *rootstock* was almost significant with p = 0.0565 and explained 22.21% of the variation (**Table 2**). *scion* explains 13.67% of variation in *B*, the spread of the early rings, and is highly significant (p = 0.000068) while *rootstock* explains more variability at 19.01% but was much less significant (p = 0.0593). For *k*, the decay, only *rootstock* was significant (p = 0.0298) explaining 22.68% of the variation. We conclude that scion modulates the widths of the first rings, as evidenced by being highly significant for modulating modeled *B* values. Rootstock, contrastingly, modulates ring widths more consistently throughout the trunk and across the lifetime of the vine by modulating the decay, *k* (**Table 2**).

**Figure 2:**
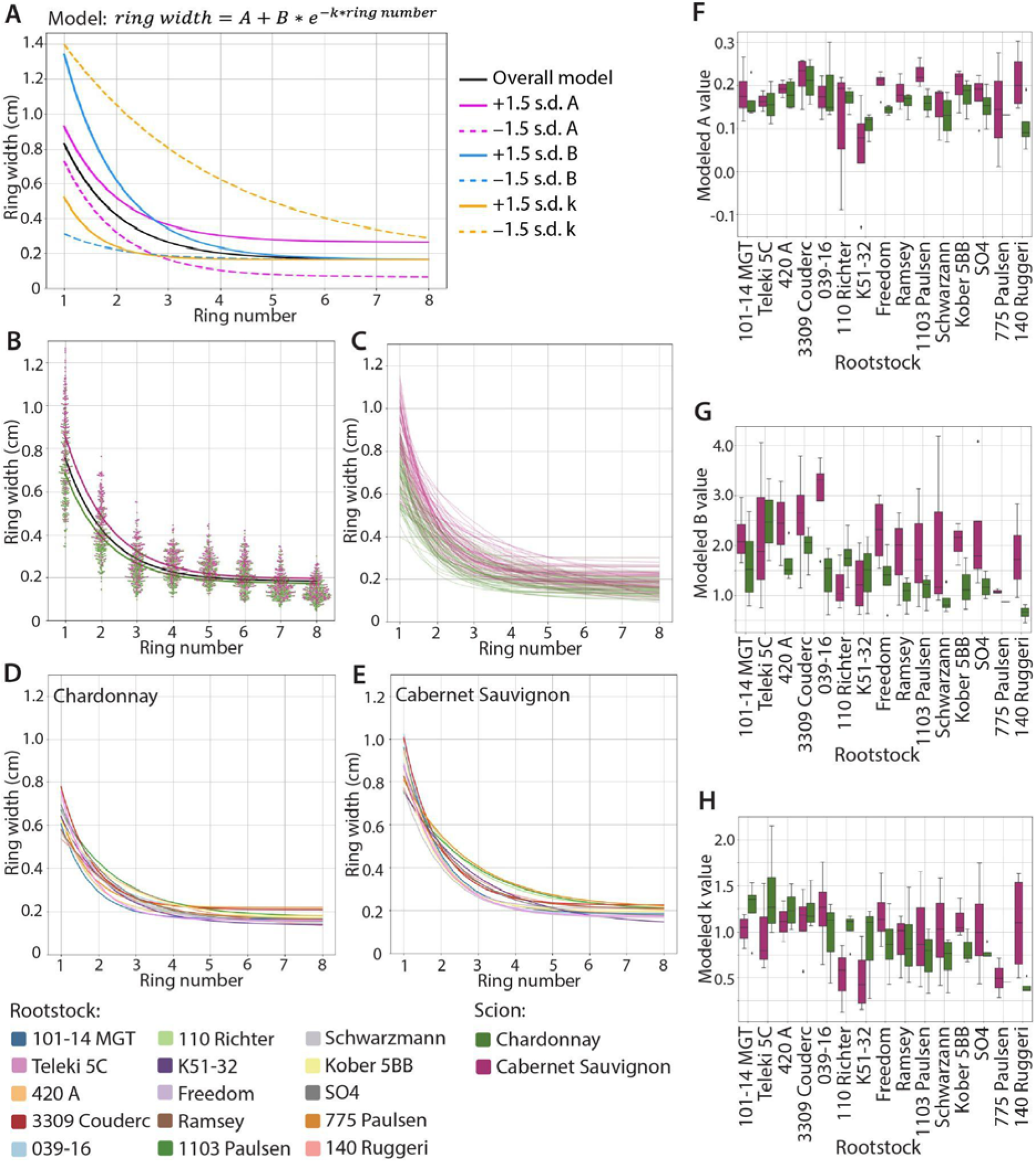
Models of ring width. **A)** For the negative exponential model *ring width* = *A* + *B*\* *ring number*, the overall model fitted to all the data (black solid line) and models fitted with values +1.5 standard deviations (solid lines) and −1.5 standard deviations (dashed lines) for *A* (magenta lines), *B* (blue lines), and *k* (orange lines). **B)** A swarmplot of all measured ring widths for Chardonnay (green) and Cabernet Sauvignon (purple) with an overall model (black) and models for each scion, **C)** models for each vine measured colored by scion, **D)** models for each rootstock for Chardonnay scions, and **E)** models for each rootstock for Cabernet Sauvignon scions. **F-H)** Boxplots of model values by rootstock and by scion for **F)** *A*, **G)** *B*, and **H)** *k.*

### Ravaz (harvest) index and juice pH are robustly correlated with scion ring and trunk width

To determine traits most correlated with ring features and if there are specific ring features that correlate with specific traits, we correlated all traits with all ring features. The ring features were modeled *A, B*, and *k* values (**Figure 2F-H**), early and late ring widths 1 to 8, outer ring widths, cambium, and total width (or, scion trunk radius). Only rings 1 to 8 were measured individually because after these, boundaries between rings and early and late wood became indistinguishable. As described in Materials and Methods and detailed in **Supporting Information Table S1**, some traits were collected only during 1995-1999, others 2017-2020, and some for both periods. To understand general trends in the data, we visualized the distribution of correlation coefficients for each trait, for each scion, across the years the trait was measured with scion trunk radius, as an overall summary (**Figure 3A**). Median Spearman’s rank correlation coefficient values between traits and scion trunk radius range from around −0.4 to 0.4. Some traits have similar correlation coefficient values between the two scions, while others are contrasting. For example, whereas pruning weight is strongly positively correlated with scion trunk radius in both scions, juice pH is similarly positively correlated in Chardonnay, but more neutral in Cabernet Sauvignon. Yield divided by pruning weight (also known as harvest, or specifically in grapevines as Ravaz, index) is strongly negatively correlated in both Chardonnay and Cabernet Sauvignon. In contrast, petiole tissue Mg and Ca traits were strongly negatively correlated with scion trunk radius in Chardonnay and more neutral in Cabernet Sauvignon (**Figure 3A**).

**Figure 3:**
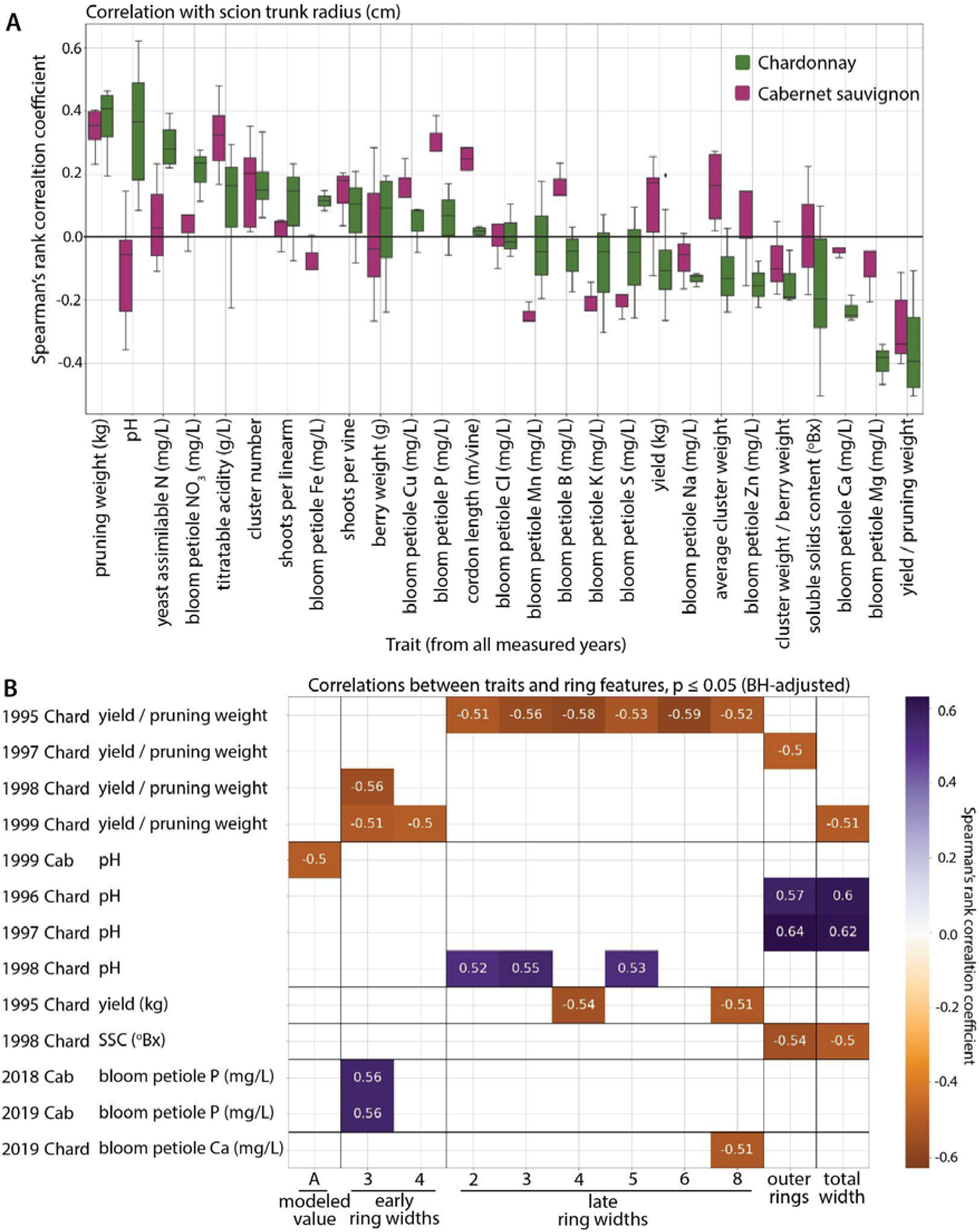
Correlations between ring features and traits. **A)** Boxplot showing distributions of Spearman’s rank correlation coefficient values for correlation between indicated traits for Chardonnay (green) and Cabernet Sauvignon (purple) and scion trunk radius (cm). **B)** 26 traits that remain significant at p < 0.05 after Benjamini-Hochberg multiple test correction. The year, scion, and trait as well as ring features (modeled values, early ring widths, late ring widths, outer ring width, or total width) are indicated. Negative (burnt orange) to positive (violet) correlation coefficient values are indicated by scale and also provided by text for each correlation.

Multiple test adjustment on 5,786 correlations between each ring feature with trait values for each scion and for each year resulted in 26 significant correlations with p ≤ 0.05 (**Figure 3B**). Our strategy in correlating each trait against each ring feature was to agnostically determine if the traits measured in years corresponding to rings (or preceding years, given the patterning of grapevine organs the year before they emerge) (Khanduja & Balasubrahmanyam, 1972; Srinivasan & Mullins, 1976; Guilpart *et al.*, 2014; Chitwood *et al.*, 2021) showed associations with each other. We did not detect any specific associations of rings with traits measured for the years they were patterned. Further, rather than building chronologies and correcting for age-related growth trends as is typically done in dendrochronology studies, we leverage the fact that all vines are the same age, allowing the widths for each ring across vines to be compared with each other. Significant correlations tended to include Chardonnay and traits measured in 1995-1999; however, this does not mean that Cabernet Sauvignon or traits measured in 2017-2020 are necessarily more weakly correlated with ring features, as these factor levels have relatively more missing data points as a result of both missing ring data (**Figure 1**) and trait data (**Supporting Information Table S1**). The missing ring data was generally a result of vines which had become scion-rooted over time and could therefore not be accurately sampled for rootstock-effects. Of the 26 significant correlations, 11 include Ravaz index and 8 juice pH (**Figure 3B**). Harvest index tended to be negatively correlated with ring and trunk widths while juice pH was positively correlated. There were no strong patterns of specific ring features correlating with traits except that there were no significant correlations with cambium width and that there were more correlations with late wood (12) compared to early wood (5). The correlation between Ravaz index and scion trunk radius was consistently negative across 1995-1999 in both Chardonnay and Cabernet Sauvignon, although the correlations were stronger and driven by variability in vines with smaller scion trunk radii in Chardonnay (**Figure 4A**). The negative correlation in Ravaz index with ring features is driven by strong positive correlations between pruning weight and ring width in both scions (**Figure 3A**). Vines with larger ring widths tended to produce more vegetative growth (higher pruning weight) while the impact on reproductive growth (yield) was weak. As a result, Ravaz index had a high negative correlation with scion trunk radius, indicating that vines with larger trunks had lower ratios of yield/pruning weight. Similar to Ravaz index, the positive correlation between juice pH and scion trunk radius is maintained in Chardonnay across 1995-1999 but is not significant for Cabernet Sauvignon (**Figure 4B**).

**Figure 4:**
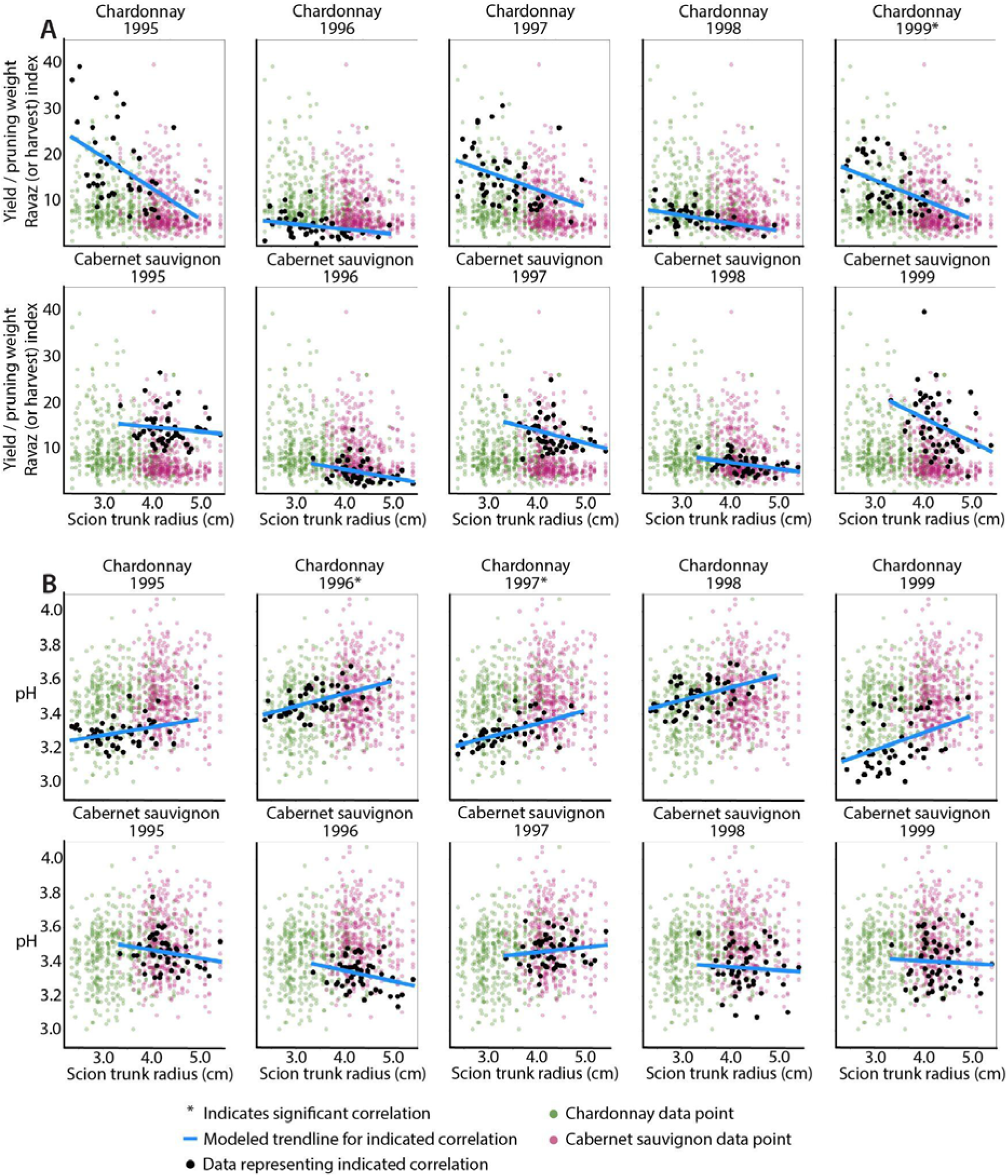
Correlations by year for harvest index and pH with scion trunk radius. Asterisks indicate significant correlations (as indicated in Figure 3B), based on multiple testing correction. For correlations between **A)** yield divided by pruning weight (harvest, or Ravaz, index) and **B)** juice pH correlations with scion trunk radius (cm) for data from each scion and indicated year (black data points) and a modeled trendline (blue) are superimposed over all data for the trait (Chardonnay, green; Cabernet Sauvignon, purple) across the years shown.

### Impact of scion trunk radius on vine physiology

To determine the physiological implications of rootstock choice, assimilation rate (A), transpiration rate (E), stomatal conductance (gsw), and leaf temperature were measured for 15 dates during the 2018-2019 growing seasons. To better synchronously measure time-intensive physiological traits with replication, three rootstocks were chosen that happen to span low to high scion trunk radius values: from low to high, Teleki 5C, Freedom, and 1103 Paulsen (**Figure 1B-C**). The repeated measures correlation coefficient (r_rm_), which fits an overall correlation coefficient value to a population of multiple measurements (Bakdash & Marusich, 2017), was used (**Figure 5**). Measurements on the three rootstocks, measured for each scion of each of the 15 dates, were used as repeated measures to calculate an overall correlation coefficient between each physiological trait and scion trunk radius. Repeated measure correlation coefficients for assimilation rate, transpiration rate, and stomatal conductance with scion trunk radius, for both Chardonnay and Cabernet Sauvignon, were positive, ranging between 0.28 and 0.37, and highly significant. The same correlations for leaf temperature, measured using an infrared temperature gun on the same leaves from which physiological traits were taken, were near 0 and not significant. For the rootstocks spanning scion trunk radius values that were measured, we conclude that vines with wider trunks are more photosynthetically active, both transpiring and assimilating more (**Figure 5**).

**Figure 5:**
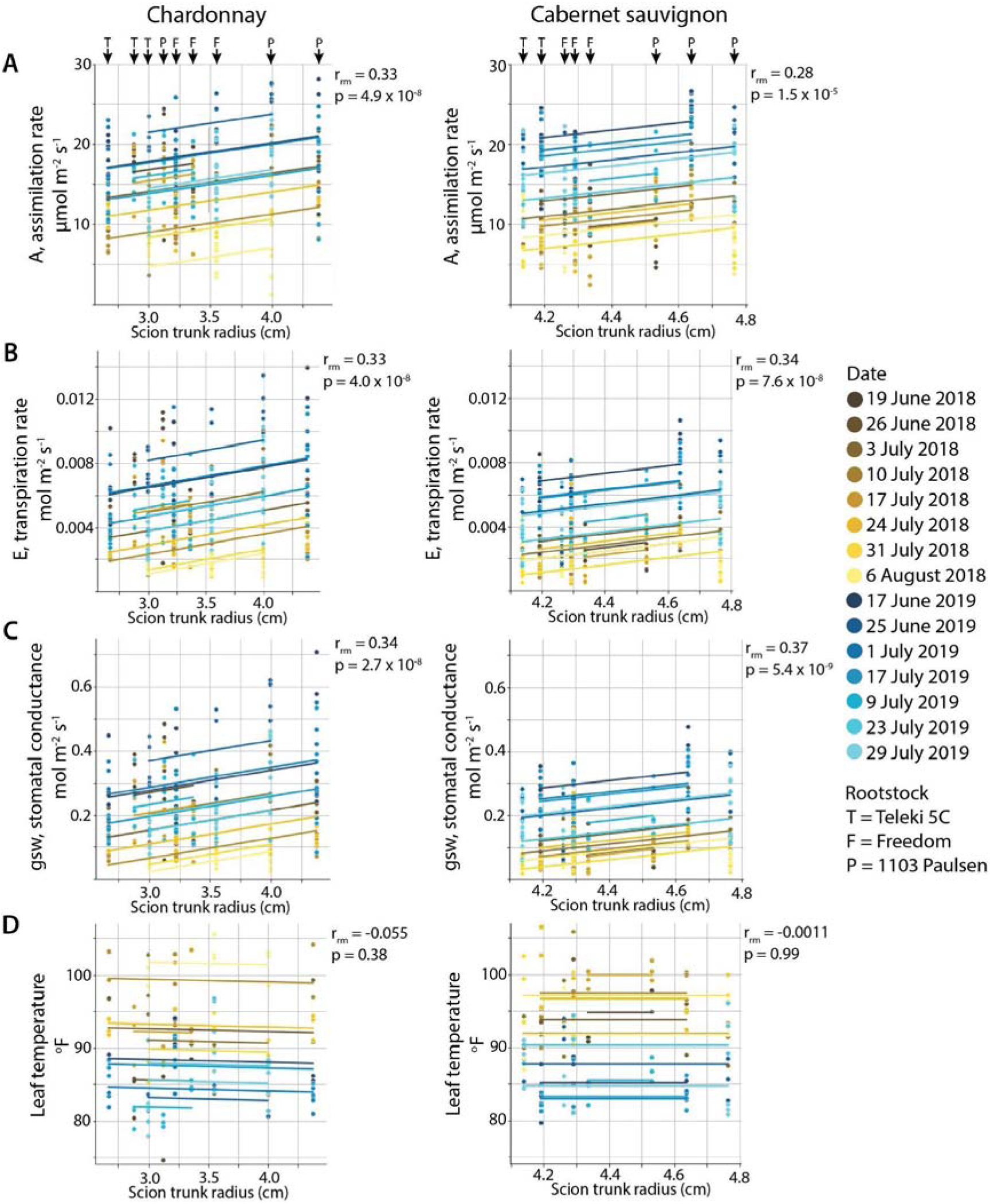
Repeated measures correlation between physiological traits and scion trunk radius for three rootstocks. Repeated measure correlation coefficient (r_rm_) and p value are shown for the overall fitted correlation between **A)** A, assimilation rate (μmol m^−2^s^−1^), **B)** E, transpiration rate (mol m^−2^s^−1^), **C)** gsw, stomatal conductance (mol m^−2^s^−1^), and **D)** leaf temperature (°F) with scion trunk radius (cm) across 3 rootstocks measured on 15 dates across two years. The scions and rootstocks (T = Teleki 5C, F = Freedom, and P = 1103 Paulsen) are indicated in the top panels and dates are indicated as shown by color, yellow shades for 2018 and blue for 2019.

## Discussion

Within the scion trunk radius ranges of Chardonnay and Cabernet Sauvignon, rootstocks modulate size in an additive, continuous fashion up to 143% (**Figure 1B-C**). These results build upon and support previous observations comparing seven year old Cabernet Sauvignon grafted onto 1103 Paulsen, 420A, and Riparia Gloire (Shtein *et al.*, 2017). Although our study does not include Riparia Gloire, which had an unusually small trunk diameter (and which was proposed as a mechanism by which hydraulic conductance and physiological performance are limited), we confirm that Cabernet Sauvignon grafted onto 1103 Paulsen has a wider scion trunk radius than 420A. Similarly, De Micco *et al.* (2018), in comparing five year old ungrafted Piedirosso to vines grafted to 420A, found that grafting limits hydraulic conductivity in desirable ways, increasing berry quality. Rather than a binary view of graft formation either inhibiting or permitting hydraulic conductance as a consequence of trunk diameter and wood anatomy, in comparing 15 rootstocks, we instead observe a continuous range of scion trunk radii. In using two scions, we also see that these effects are additive, layered on top of the larger scion effect, mostly preserving the ranking of scion trunk radii conferred by different rootstocks (**Figure 1**; **Table 2**).

Scion trunk radius is the summation of annual ring widths and by measuring the outcome of 30 years of growth, we are able to identify the mechanisms by which rootstocks alter wood anatomy (**Figure 2**). Using the model *ring width* = *A* + *B* * *e*-^*k*\**ring number*^, scion most strongly modulates *B*, the spread of the first rings (**Figure 2A; Table 2**). Considering that scions were whip grafted to rootstocks in their first year of growth, scion effects are expected to dominate in the absence of any rootstock. Rootstock most strongly modulates the decay of ring width *k* thereafter, which can be interpreted as a consistent effect across all rings for the lifetime of the grafted vine. How rootstocks modulate annual ring width is illuminated by previous studies demonstrating how the environment does so. In grapevines and other lianas, rings arise from high cambial activity early in the growing season and less at the end, creating the alternating seasonal pattern of early and late wood (Berstein & Fahn, 1960; Pratt, 1974; Kozlowski, 1983; Ewers *et al.*, 1990; Wheeler & LaPasha, 1994). As shown previously by Munitz *et al.* (2018), ring width increases with water availability, especially in the early growing season (late deficit) when cambial activity is greatest. Hypotheses of rootstocks (Shtein *et al.*, 2017) and grafting (De Micco *et al.*, 2018) altering physiological performance by restricting hydraulic conductance are derived from extrapolating from these well-studied environmental effects of water availability in the early season coinciding with meristem activity and the development of wood anatomy.

The impact of environment and rootstocks on physiological performance are derived both from ring width and trunk diameter as well as parallel changes in vessel anatomy affecting overall hydraulic conductivity of the vine. Generally in vines vessel length and diameter are correlated with stem diameter (Ewers & Fisher, 1989; Jacobsen *et al.*, 2012, 2015) and grafting impacts vessel frequency and size in peach, cherry, and apple as well (Olmstead *et al.*, 2006; Gonçalves *et al.*, 2007; Tombesi *et al.*, 2009). Density of vessels is not affected by irrigation, but overall vessel diameter and diameter of large vessels increases with water availability, especially under late deficit in the early growing season (Munitz *et al.*, 2018; Netzer *et al.*, 2019). A previous study noted increases in the frequency of more narrow vessels in late wood in response to grafting (De Micco *et al.*, 2018). However, when considered with annual ring width and the distribution of vessel diameters, grafting reduces hydraulic conductivity (De Micco *et al.*, 2018) and rootstocks with larger trunk diameters increase it (Shtein *et al.*, 2017). Vessels are visible in our X-ray CT images and can be measured (**Figure 1**), but as the pixel resolution is limited to 63.9 μm, we are unable to measure narrow vessel diameters that constitute an important segment of vessels in the bimodal distribution of grapevine (Munitz *et al.*, 2018) and which grafting is reported to increase in frequency (De Micco *et al.*, 2018). The impacts of rootstocks on hydraulic connectivity can nonetheless be inferred from physiological performance, as photosynthetic assimilation rate, transpiration rate, and stomatal conductance all decrease when water status is impaired (Romero & Martinez-Cutillas, 2012). Across these three rootstocks, we observe highly significant correlations between physiological parameters known to be affected by hydraulic conductivity with scion trunk radius (**Figure 5**). Although the extent of alignment between rootstock and scion was not evaluated in this study, vines with complete alignment may have higher levels of transpiration than those with partial alignment, indicating that the success of grafting itself may also play a role in physiological differences. However, in previous work this did not correspond to hydraulic conductivity differences across the graft measured in three year old vines at the end of the study (Marín *et al.*, 2022).

Vegetative growth can indicate water availability (Tyree & Ewers, 1991; Munitz *et al.*, 2017), and from this perspective it is not surprising that some of the most correlated traits with rootstock-induced changes in scion trunk radii are related to growth (**Figures 3-5**). Indeed, a previous study of this vineyard suggested that the 1998 reduction in yield observed may be due to a dry 1997 dormant season (Migicovsky *et al.*, 2021). The cumulative precipitation for 1997 was 2,533 inches. In comparison, precipitation for 1995, 1996, 1998, and 1999, ranged from 3,377 to 7,775 inches, indicating that there was substantially more rainfall during these years (Migicovsky *et al.*, 2021). The 1998 season is one of the same years where the correlation with Ravaz index, the ratio of yield to pruning weight, is weak for both Chardonnay and Cabernet Sauvignon (**Figure 4A**). However, overall, the correlations with Ravaz index are some of the strongest. As scion trunk radius increases in Chardonnay, Ravaz index decreases, driven mostly by year-to-year variation in vines with narrow trunk diameters from 1995 to 1999 (**Figure 4A**). The relationship is less strong for Cabernet Sauvignon. Similar to 1998, the correlation with the Ravaz index from 1996 is also poor. Both 1996 and 1998 were years previously reported to have low yields, and as a result, low Ravaz indexes. Although the reason for low yields in 1996 is unclear, among the 1995-1999 data, it was the year with the highest pruning weights, indicating a stronger investment in vegetative growth (Migicovsky *et al.*, 2021). Underlying the strongest correlations between Ravaz index and scion trunk radius observed in both scions are years of higher yields and lower pruning weights (1995, 1997, and 1999) (**Figure 3A**), thus indicating that in years that enable high reproductive growth, yields are lower in large scions. While there was a modestly positive correlation between yield and scion trunk radius in Cabernet Sauvignon, the correlation for Chardonnay was modestly negative which likely explains why the correlation between Ravaz index and scion trunk radius was higher for Chardonnay. Taken together, these findings indicate that generally vines with a larger trunk radius will have a lower Ravaz index as a result of having higher vegetative growth (pruning weight). This relationship is stronger in Chardonnay, which also generally had smaller vines (**Figure 1B**) as well as lower yields in comparison to Cabernet Sauvignon (Migicovsky *et al.*, 2021). A lower Ravaz index is not desirable for grape growers, because it leads to higher management costs relative to the increase in profit (or yield).

Similar to pruning weight, juice pH had a strong positive correlation with scion trunk radius, but only in Chardonnay, which was consistently observed across 1995 to 1999 (**Figures 3, 4**). Juice pH is potentially influenced by potassium uptake of rootstocks (Ruhl, 1989; Kodur, 2011), but we did not observe strong correlations between petiole potassium levels and scion trunk radius (**Figure 3A**). We mention correlations with juice pH because it is strongly positive and scion-specific, and it is consistent across years (**Figure 4B**), in addition to showing that effects on hydraulic conductivity and growth associated with ring width and trunk diameter can potentially affect berry quality through indirect mechanisms of water and nutrient uptake (Keller *et al.*, 2012; Mantilla *et al.*, 2018; Migicovsky *et al.*, 2019). We focus on the correlation of traits with scion trunk radius as a cumulative indicator of rootstock-mediated effects on scion annual ring widths, but in correlating with each ring feature, we note that there are only 5 significant correlations with early wood ring widths compared to 12 significant correlations with late wood ring widths (**Figure 3B**). De Micco *et al.* (2018) also note that grafting induces more, narrower vessels creating wider latewood rings compared to ungrafted vines. Although speculation, it might be that water availability in the early season environment corresponding with cambial activity mostly affects early wood ring width (Munitz *et al.*, 2017; Netzer *et al.*, 2019) whereas rootstocks act relatively more on latewood and narrower vessels (**Figure 3B**; (De Micco *et al.*, 2018)), contributing to a consistent genetic, environmentally-independent mechanism of modulating wood anatomy.

Regardless of mechanism, our results show that rootstocks act consistently over decades to modulate growth and other scion traits. Although the significant correlations after multiple test adjustments are biased by small amounts of missing data (**Figure 3B**; **Supporting Information Table S1**), the strength, direction, and scion-specificity of the correlations are consistent for traits from the beginning (1995-1999) to the end (2017-2020) of the 30 year life of the vineyard we studied (**Figure 3A**). Vessels in *V. vinifera* are only active 1-3 years before inactivation and no vessels are active after 7 years (Pratt, 1974; Tibbetts & Ewers, 2000; Pratt & Jacobsen, 2018). From this perspective, the continuous range of scion trunk radii conferred by grapevine rootstocks (**Figure 1**) is a symptom and consequence of consistent modulation of wood anatomy across the years (**Figure 2**) that renews the hydraulic effects on rootstocks on viticultural traits (**Figures 3-4**) through physiological mechanisms (**Figure 5**). Just as used in dendrochronology to infer climatic conditions from secondary growth, annual rings can be used as a way to allow the vines themselves to report on genetic and environmental effects that are modulating their performance. The wide-ranging and continuous effects of rootstocks on scion wood anatomy are a powerful way that grape growers can precisely modulate the vegetative growth versus yield, and indirectly berry and resulting wine quality, by altering hydraulic conductivity consistently over years, with important implications for the use of rootstocks in all woody perennial species.

## Acknowledgements

We gratefully acknowledge all the individuals involved in maintaining the vineyard evaluated for this study, in particular Ernie and Jeff Dosio (Pacific Agrilands Inc., Modesto, CA). We also thank Alex Freeman (Michigan State University) for early contributions to data analysis.

## Author Contribution

ZM, JFS, JLH, DRH, LMJ, MK, RKS, and PC collected vineyard trait data and/or prepared wood for analysis. MYG and JM collected and prepared X-ray CT images. DHC measured ring widths. TA, ARA, JDD, BMG, IK, MRM, ELM, AWS, ALD, SS, and DHC analyzed data. ZM and DHC coordinated research, data analysis, and manuscript writing. DHC wrote a first draft of the manuscript which all authors read, commented on, and edited.

## Data Availability

X-ray CT cross-sections with landmarks are deposited on Dryad: http://dx.doi.org/10.5061/dryad.gqnk98sqf. All data and code to reproduce results are posted on the Github repository https://github.com/DanChitwood/grapevine_rings.

**Supporting Information Table S1:** Numbers of measured samples for each trait, for each scion, for each year.

